# Reaction kinetics of procainamide dye derivatization of N-linked glycans to enable robust process analytical workflows for glycoprotein-based biologics manufacturing

**DOI:** 10.1101/2025.10.15.682640

**Authors:** Nurulhuda Binte Suaini, Aditya Narvekar, Shishir P.S. Chundawat

**Author notes:** Both authors contributed equally to this work (Nurulhuda Binte Suaini, Aditya Narvekar).

## Abstract

N-glycosylation is a post-translational modification of proteins that represents a critical quality attribute (CQA) for therapeutics like monoclonal antibodies (mAbs), directly affecting drug efficacy, safety, and stability. Real-time CQA monitoring analytical platforms depend on rapid N-glycan release and fluorophore labeling chemistries to support automated bioprocess analytics during mAb manufacturing. Procainamide is a well-known fluorophore used for released N-glycans reducing sugar aldehydes labeling that offers both high fluorescence and mass spectrometry detection sensitivity comparable to several commercial reagents available in the market. However, currently there are no studies that optimize its use and long incubation times are often reported in the literature for procainamide labeling of N-glycans that has limited its use in time-sensitive workflows relevant to various stakeholders in industry, academia, and regulatory agencies. Here, we have systematically determined the combined procainamide labeling via reductive amination/reduction reaction kinetics at various incubation intervals, ranging from 1 min to 12 h, using N-glycans isolated from model biologic glycoprotein trastuzumab (TmAb). Labeling efficiencies were quantified using high-performance liquid chromatography with fluorescence detection (HPLC-FLR), and detailed reaction parameters were determined by fitting suitable kinetic models. Results indicate that most N-glycans reached over 95% labeling efficiency within 1 hour at the desired reaction temperature. Interestingly, N-glycan structural features, particularly galactosylation and fucosylation levels, significantly influenced the labeling reaction rate. Fucosylated glycans exhibited up to 4-fold higher reaction rate constants than non-fucosylated forms, whereas increased galactosylation levels was associated with slower reaction rate. These results provide essential kinetic benchmarks for incorporating procainamide labeling for released N-glycans, and facilitating more efficient analytical workflows for Process Analytical Technology (PAT) focused on biologics N-glycan analysis in both research and industrial settings.

## Introduction

N-glycans are complex carbohydrates often attached as glycoconjugates to other biomolecules (e.g., proteins, RNA) that play essential roles in biological systems, influencing protein folding, stability, and cell signaling. In therapeutics like monoclonal antibodies (mAbs), N-glycosylation is a critical quality attribute (CQA) that modulates pharmacokinetics, pharmacodynamics, and immune effector functions through modification at the conserved Asn297 residue on the heavy chain.^1, 2^ Aberrant glycosylation patterns are also associated with various disease states, including cancer, where glycans can serve as diagnostic or prognostic biomarkers.^1,3^ Consequently, consistent monitoring of N-glycosylation composition during biologics manufacturing is crucial to ensure the safety and efficacy of biological drugs.

Process analytical technology (PAT) platforms, such as the N-GLYcanyzer workflow developed by our team,^4, 5^ have been developed to facilitate automated, real-time monitoring of mAb glycosylation during upstream cell culture biomanufacturing. The N-GLYcanyzer workflow integrates automated sampling from bioreactors, sample preparation to isolate mAbs/glycans, and high-performance liquid chromatography (HPLC) to provide detailed glycoform composition and antibody titer data during the entire upstream process durations dictated majorly by the fluorescent dye and labeling chemistry used.^4, 5^ Fluorescent labeling is a key step in this workflow because native released N-glycans lack chromophores or fluorophores, making direct detection of released native N-glycans with high sensitivity very challenging.^6^ Chemical derivatization with fluorescent dyes, such as 2-aminobenzamide (2-AB), procainamide, or instant procainamide or 2-(diethylamino) ethyl-4-({[(2,5- dioxopyrrolidin-1-yl)oxy]carbonyl}amino)benzoate dye such as Agilent’s InstantPC™ (IPC), that enhance glycan detection sensitivity and enables quantitative glycan analysis by fluorescence and/or mass spectrometry (MS).^7, 8^

Labeling of N-glycans for HPLC analysis often proceeds typically via a reductive amination, in which the aldehyde group at the reducing terminus of the enzymatically released N-glycan reacts with an amine-containing fluorophore dye to form a Schiff base that is then subsequently reduced to a stable secondary amine using a suitable reducing agent.^8, 9^ This approach provides a stoichiometric, covalent label attached per glycan, to enable reliable quantification and structural elucidation of all major N-glycans found in biologics. The reaction mechanism for the procainamide fluorophore reductive amination step is shown schematically in **Figure 1**.^9-11^

**Figure 1.**
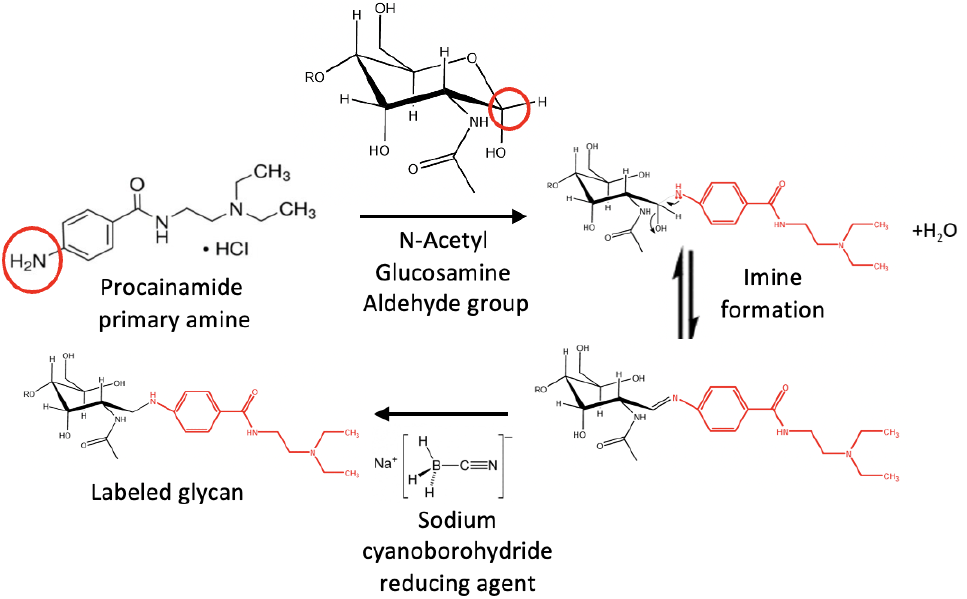
Reaction mechanism for procainamide labeling of N-glycans via reductive amination chemistry. The reducing end of the enzymatically released N-glycans reacts with the primary amine of procainamide to reversibly form a imine (Schiff base), which is subsequently reduced by sodium cyanoborohydride (NaBH□CN) to yield a stable secondary-amine labeled glycan.

Among available reductive amination dyes for glycan labeling, 2-AB remains the benchmark for most reported Hydrophilic Interaction Liquid Chromatography (HILIC) based labeled glycan analysis due to its reproducibility and cost-effectiveness. However, its limited ionization efficiency restricts its application in MS workflows.^6, 8^ To overcome these limitations, the InstantPC (IPC) label was introduced, enabling rapid (≤5 min) glycan derivatization through NHS-carbamate chemistry and improved sensitivity of labeled IPC fluorophore. Despite its strong fluorescence and MS response, labeling dyes like IPC are expensive and this can limit its use in high reagent volume use applications like real-time bioprocess monitoring.^7, 12-15^

Procainamide has emerged as a practical alternative, combining high fluorescence intensity, superior MS ionization efficiency (up to 30-fold greater than 2-AB), and comparable chromatographic behavior of labeled glycans for HILIC analysis. It employs a similar well-established reductive amination chemistry, allowing straightforward integration into existing analytical workflows.^6,8^ As PAT platforms evolve to include MS-based monitoring, procainamide is an attractive candidate for real-time glycan analysis deploying automated sampling/labeling tools such as N-GLYcanyzer.^4, 16^ However, one major issue is the poor reaction kinetics understanding and hence prolonged labeling time (∼3 h at 60 °C) often reported for procainamide unlike IPC, which can limit its adoption as a reagent for glycoform analysis.

This study aims to characterize procainamide labeling kinetics in detail for optimizing real-time released N-glycan monitoring applications of relevance to the pharmaceutical industry. A time-course study was conducted to evaluate labeling efficiency across multiple incubation times (1 min–12 h) using N-glycans isolated from trastuzumab (TmAb) produced in Chinese Hamster Ovary (CHO) cells. Labeling efficiency was quantified by HPLC-fluorescence, and kinetic modeling was applied to describe the reaction kinetics. This work establishes the foundation for accelerated PAT-compatible N-glycan labeling workflows that support real-time monitoring of mAb glycosylation in biopharmaceutical manufacturing using procainamide as the labeling reagent.

## Materials and methods

### Reagents

PNGase F was purchased from Bulldog-bio Inc., Portsmouth, NH, United States. Procainamide hydrochloride, N-lauroylsarcosine, Dithiothreitol, Sodium cyanoborohydride, Ammonium formate was purchased from Sigma-Aldrich, Inc. St. Louis, MO, United States. Acetonitrile, Glacial acetic acid was purchased from VWR Chemicals, Radnor, PA, United States. Dimethyl sulfoxide and Octanal was purchased from Millipore Sigma, Burlington, MA, United States.

### Fed batch bioreactor production of TmAb and its purification

Trastuzumab biosimilar was produced using fed-batch bioreactor set up and characterized as described previously.^4^ Briefly, after harvesting the fed batch reactor after ∼14 days of cell culture, the TmAb in the supernatant was purified as per manual instructions using rProtein A GraviTrap column (Cytiva, Wilmington, DE) and the purified TmAb concentration was estimated on HPLC using a Bio-Monolith protein A column (Agilent Technologies, Santa Clara, CA).

### Protein denaturation and N-glycans release

The purified Trastuzumab (>50 µg) sample was mixed with a protein denaturant solution containing denaturant (N-lauroylsarcosine) and a reducing agent (dithiothreitol). The protein was denatured using a stock of 10% (w/v) N-lauroylsarcosine (N-LS) and 400 mM dithiothreitol (DTT) at the final working concentration of 0.5% and 20 mM, respectively. Next, 2 µL of PNGase F was added to catalyze the enzymatic release of N-glycans. The solution was mixed thoroughly to ensure homogeneity. The sample was incubated at 50□°C for 5 minutes to denature the protein and facilitate rapid glycans release.

### Procainamide labeling and cleanup of released N-glycans

Procainamide labeling solution was prepared using 70% dimethyl sulfoxide (DMSO) and 30% glacial acetic acid (AcOH) as the solvent mixture. This solvent mixture was used to dissolve procainamide hydrochloride (11 mg/100 μL) and sodium cyanoborohydride (NaBH□CN) (6 mg/111 μL), and was mixed thoroughly to ensure complete dissolution. For complete solubilization of the reducing agent, DI water was added (30 μL/6mg NaBH□CN). The resulting solution was used as the labeling reagent. The denatured mAb and glycan released samples was kept at room temperature for 2 minutes after the PNGase reaction step. Procainamide labeling solution was added at a 1:1 volume ratio relative to the sample. The tubes were capped and incubated at 65□°C until the respective sampling time points to ensure complete labeling by reductive amination. Labeling reactions were stopped at thirteen defined time points: 1□min, 5□min, 10□min, 15□min, 30□min, 1□h, 1.5□h, 2□h, 2.5□h, 3□h, 3.5 h, 6□h, and 12□h. Each time point sample was prepared in duplicates, resulting in two independently labeled samples per reaction time interval.

After incubation, post-labeling clean-up was carried out using an octanal-based liquid-liquid extraction protocol. Briefly, the labeled sample was mixed with octanal in a 1:2 volume ratio (sample: octanal) and vortexed for 30 seconds. The organic and aqueous phases were separated by centrifugation for 1 minute at 3000 rpm using microcentrifuge (Minispin, Eppendorf, Hamburg, Germany). The upper organic layer was removed, and the clean-up step was repeated two more times to remove excess labeling dye. After the final clean-up step, the lower aqueous phase containing labeled glycans was collected and transferred to an HPLC vial for analysis.

### HPLC analysis of labeled glycans

Procainamide-labeled glycans were analyzed using an Agilent Online Monitoring 1290 HPLC (Agilent Technologies Inc., Germany) system with a fluorescence detector. The detector was set to an excitation wavelength of 308 nm and an emission wavelength of 359 nm. System operation, chromatographic data acquisition, and peak analysis were performed using Agilent OpenLab CDS software. Chromatographic separation was achieved with an AdvanceBio Glycan Mapping column (120 Å, 2.1 × 150 mm, 2.7 µm particle size; Agilent Technologies Inc.). The mobile phase A consisted of 100 mM ammonium formate buffer (pH ∼4.4), and acetonitrile (ACN) was used as mobile phase B. The ammonium formate was delivered in a linear gradient from 15% to 100% over 48 minutes at a flow rate of 0.5 mL/min. To maintain stability, the samples were stored in the HPLC multisampler compartment at 10°C before injection, and the injection volume was set to 1 µL. Each procainamide-labeled glycan sample was injected twice to confirm HPLC analytical reproducibility. Two experimental replicates were carried out and thus, there were 4 data points were collected for each sample condition.

### Data processing and kinetic curve fitting

Agilent OpenLab Chromatography Data System (CDS) software provides default quantitative integrations methods to estimate area under the curve for analyzed glycans. The lower intensity glycan peaks which were not detected using the default integration methods were manually integrated. Standard immunoglobulin G (IgG) glycoforms were also identified by matching their chromatography peaks to specific retention times of known standards. The fluorescence signal area of each glycan peak was integrated to quantify labeling intensity. These data were exported for analysis and development of a kinetic model.

Three different plots were generated to illustrate the kinetic modeling of the time-series incubation study. A comparative overview of the plot was created by overlaying the experimental data. This was achieved by combining the plots of all nine N-glycans, which involved averaging the fluorescence area intensities across duplicates for each glycan. In addition to this composite view, individual kinetic plots were generated separately for major glycans (i.e., G0F, G1F, G1F′, G2F) and minor glycans (i.e., G0, G1, G1′, G0F-N, Man5), each based on raw area intensities from duplicate injections. Kinetic modeling was completed using Python 3.10 on Google Colab, with AI-assisted tools and custom scripts developed with the SciPy and NumPy libraries. Three separate kinetic plots were fitted to an exponential saturation model as shown below to allow a simple interpretation of the predicted reaction rates.

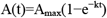

Where A(t) is the fluorescence intensity at time t, A_max_ represents the maximum signal, and k is the rate constant (h□^1^).

Nonlinear least squares fitting used scipy.optimize.curve_fit, and the goodness of fit was evaluated with the coefficient of determination (R^2^). Quantitative markers such as kinetic parameters, k and A_max_ were estimated from the exponential saturation model fits to evaluate labeling reaction rates and relative signal intensities across different glycan datasets. These values were further analysed to examine the relationship between glycan structural features, core fucosylation, reducing-end accessibility, and its effects on labeling efficiency.

## Results and Discussion

### Trastuzumab biosimilar released N-glycans structural and HPLC analysis

Previous studies have demonstrated that the glycan structure, particularly the composition of terminal sugar residues such as mannose, sialic acid, fucose, and galactose, can influence the efficacy of therapeutic immunoglobulin (IgG) mAbs.^17^ The nine procainamide-labeled N-glycans analyzed in this study for the model Trastuzumab biosimilar are common IgG Fc glycoforms typically produced by mAbs expressed in CHO cells. These include both complex-type and high-mannose structures, which differ in their branching patterns, terminal sugars, and other core modifications. **Supplementary Table S1** shows the different glycan structures analyzed in this study, while **Figure S1** and **Table S2** depict representative examples of a chromatogram depicting elution profiles of all glycans analyzed from the Trastuzumab biosimilar along with the quantification of the peak areas under the curve. As a biantennary complex glycan, G0 ends in N-acetylglucosamine (GlcNAc) on both antennae rather than terminal galactose residues. The innermost GlcNAc of its fucosylated analogue, G0F, has an additional α-1,6-linked fucose attached. This alteration is common in IgG antibodies and has been shown to affect glycan shape and Fc receptor binding.^18^ The monogalactosylated glycans G1 contain an α-1,6-arm, while its structural isomer, G1′, features galactose on the α-1,3-arm. The fucosylated variants of both forms, G1F and G1F′, are also found to contain core α-1,6-fucose while maintaining the same galactosylation positions. G2F is the fully galactosylated biantennary glycan with a core fucose, and both antennae are terminated with galactose.^19, 20^ In a structurally different form, G0F-N, the core mannose has a bisecting β1,4-linked GlcNAc residue attached, which is known to reduce glycan core flexibility and may limit access to enzymes or labeling reagents.^2^ Compared to other complex-type glycans, Man5 is the only open, unbranched structure with five mannose residues branching from a chitobiose core (Man□GlcNAc□). The absence of a terminal galactose or GlcNAc moiety increases its solvent accessibility and may affect labeling efficiency as well.^21, 22^

Variations in the glycan structures have been studied for their impact on pharmacokinetics and pharmacodynamics. For example, the presence of terminal galactose residues increases complement-dependent cytotoxicity (CDC) antibody binding to C1q, therefore enhancing complement activation and facilitating the killing of target cells.^14^ Beyond their biological effects, previous studies indicate that the structure, size, and charge of the glycans could potentially influence labeling efficiency. Labeled and unlabeled glycans have been reported to result from partial degradation or incomplete derivatization during the labeling process and sample preparation, such as desialylation or defucosylation.^23, 24^ However, the underlying mechanisms of how and why glycan structural features influence labeling kinetics, especially the rates of derivatization during reductive amination, has remained underexplored. Here, we will discuss our findings that provides new insights for future research on the relationship between glycan structural features and labeling efficiency using reductive amination chemistry.

### Estimation of kinetic parameters of Trastuzumab glycan procainamide labeling

Here we investigated the labeling efficiency of procainamide in a time incubation series experiment involving nine N-glycans. The glycan fluorescent labeling data were fitted to an exponential saturation model as depicted in **Figure 2** to estimate the kinetic rate constant (k) and the maximum fluorescence signal (A_max_) for each glycan (shown in **Table 1**). The exponential saturation model best fits the kinetics of procainamide labeling through the reductive amination reaction.^25-27^ After enzymatic cleavage, the released glycan’s reducing end exists in equilibrium between the cyclic hemiacetal form and the open-chain aldehyde.

**Table 1.** Exponential saturation model fitted parameters and uncertainties (A_max_, k, and R^2^) for all major and minor N-glycans found in TmAb based on averaged fluorescence data (n = 4; duplicate reactions × duplicate injections).

| Glycans | $A_{\max}$ | $k$ | $R^2$ |
| --- | --- | --- | --- |
| G0 | 16.89±0.22 | 2.40±0.06 | 0.81 |
| G0F | 186.09±0.09 | 8.68±0.10 | 0.88 |
| G1 | 5.53±0.08 | 3.16±0.08 | 0.80 |
| G1' | 2.13±0.01 | 0.80±0.03 | 0.75 |
| G1F | 76.39±0.20 | 3.70±0.10 | 0.88 |
| G1F' | 39.05±0.14 | 3.28±0.04 | 0.74 |
| G2F | 11.05±0.08 | 2.18±0.06 | 0.86 |
| G0F-N | 5.75±0.04 | 5.06±0.02 | 0.74 |

**Figure 2.**
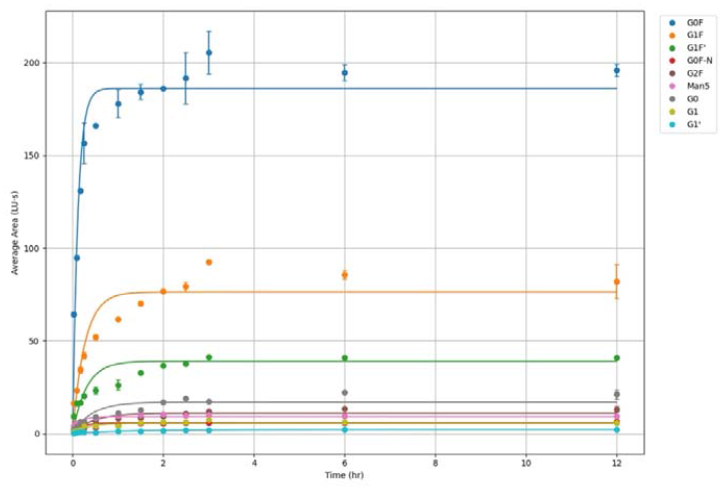
Exponential saturation model fitted for procainamide labeling reaction kinetics for nine N-glycans relevant to TmAb. Each data point represents the mean (filled circle) ± standard deviation (error bars) range of four replicate measurements obtained from two independent labeling reactions, each analyzed in duplicate HPLC-FLR injections. The fitted curves illustrate the time-dependent increase in fluorescence area under the curve (AUC) with increasing reductive amination reaction time.

The primary amine of procainamide reacts with the aldehyde group to form a reversible imine intermediate. In aqueous conditions, imine can readily hydrolyse back to aldehyde and amine, but an excess of procainamide can shift this equilibrium toward imine formation. The reducing agent, sodium cyanoborohydride, can then irreversibly reduce the intermediate imine to a stable secondary amine labeled glycan. Since procainamide and NaBH□CN are present in excess relative to the glycan substrate, their concentrations remain essentially constant during incubation. Under these conditions, the reaction can be assumed to follow a pseudo-first-order reaction rate model, where the labeling rate is dependent only on the glycan concentration. The modelling analysis shows that the fluorescent signal (area) is proportional to the concentration of labeled glycan, increasing rapidly before plateauing as the substrate availability is reduced.^15, 28^

Figure 2. provides a comparative overview of the reaction kinetics of nine N-glycans, showing the rate and extent to which each glycan is labeled with procainamide over time. Each curve represents an exponential saturation model fitted to the average fluorescence signal over time, along with its kinetic parameters such as k and A_max_. Glycans like G0F (k = 8.68 h□^1^) and Man5 (k = 6.44 h□^1^) showed rapid labeling, reaching near-maximum saturation signals within the first hour of initiating the reaction. In contrast, G1′ labelled much more slowly (k = 0.80 h□^1^), likely due to structural effects that restrict access of reagents to the reducing aldehyde end. Although G1 and G1’ have identical monosaccharide compositions, the arm-specific position of galactose may cause subtle conformational differences, limiting reagent access to the reducing aldehyde end during chemical derivatization. Glycan isomerism can influence its stereochemistry and structural group orientation, which may impact the chemical reactivity of the aldehyde group. The overall kinetics dataset showed an acceptable fit to the simple exponential saturation model despite minor deviations at specific time points.^29^

### Reductive amination reaction profile and rate parameters of major and minor N-glycans

**Figures 3** and **4** show the individual plots of major and minor glycans, respectively. Major glycan G0F lacking the galactose residues, showed the highest labeling intensity of 186.1 LU-s, making it the most efficiently labeled glycan. This may result from minimal steric hindrance near the reducing end, where procainamide labeling occurs. G1F, galactosylated on the α-1,3 arm was the next most abundant glycan, with 76.39 LU-s, consistent with efficient labeling. Its isomer G1F’ differs in conformation and showed a lower A_max_ and k, indicating that small structural changes can easily affect reducing end accessibility. G2F, which is galactosylated on both arms, was the most sterically hindered and exhibited a slow labeling efficiency of 2.18 hr□^1^. The trend for major glycans indicates that increased galactosylation is correlated with decreased labeling efficiency.^9, 30, 31^ The minor glycan Man5 shows a linear, branched structure which appears to be a key factor in faster kinetics, as it achieved the fastest labeling rate of 6.44 h□^1^ and reached saturation quickly. Its highly compact structure and greatly reduced steric hindrance likely allows for more efficient access by procainamide label.^8^ Other minor glycans, like G0, G1, and G1’, composed of biantennary branches or containing a galactose residue, showed slower labeling kinetics likely due to increased steric crowding. These findings demonstrate that glycan architecture is a crucial factor in the kinetics of reductive amination. The observed reaction dynamics reveals how branching patterns and terminal sugar composition influence N-glycans labeling efficiency, emphasizing the broader connection between released N-glycan structure and its chemical reactivity at the reducing end.^32, 33^

**Figure 3.**
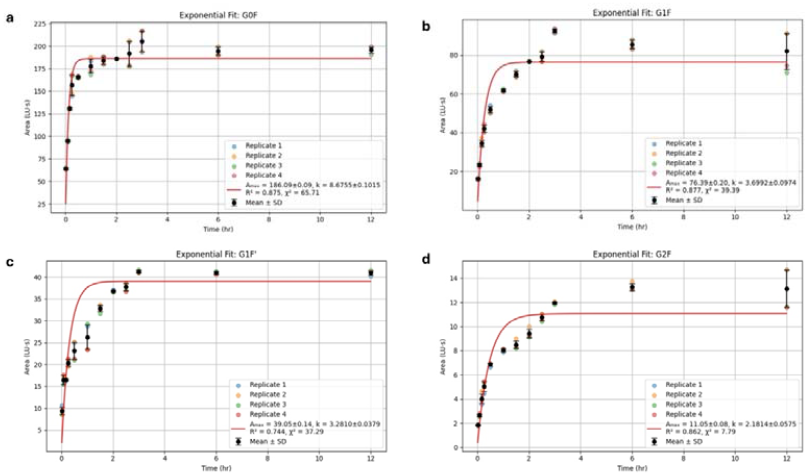
Individual time-course profiles and exponential saturation model fits for major glycans - G0F(a), G1F(b), G1F′(c), and G2F(d). Data represents mean fluorescence areas (AUC) from four replicate measurements (duplicate labeling reactions, each injected twice on HPLC-FLR). Here means are shown as filled circles with standard deviation for error bars.

**Figure 4.**
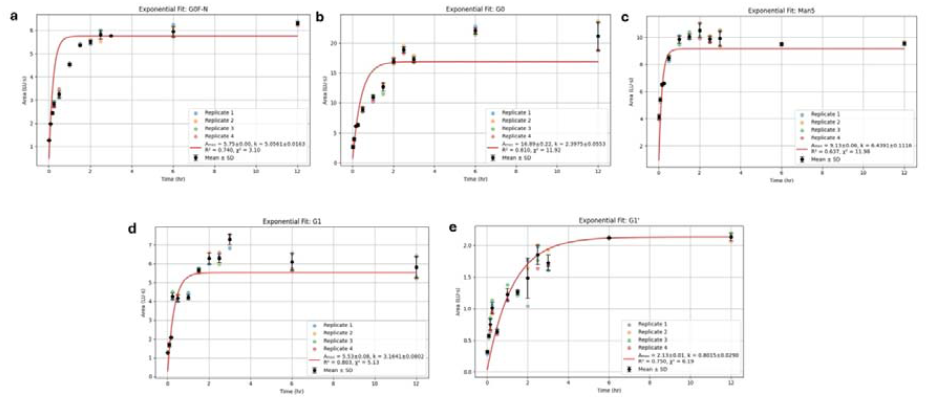
Individual time-course profiles and exponential saturation model fits for minor glycans – G0F-N(a), G0(b), Man5(c), G1(d), and G1′(e). Data represent mean fluorescence areas (AUC) from four replicate measurements (duplicate labeling reactions, each injected twice on HPLC-FLR). Here means are shown as filled circles with standard deviation for error bars.

### Impact of fucosylation of N-glycans core structure on reductive amination reaction

Based on the released glycans’ reductive amination reaction kinetic profile, it was observed that fucosylated glycans exhibited faster labeling efficiency, consistently yielding a higher rate constant as compared to non-fucosylated, also known as afucosylated glycans, as summarized in **Table 2**.

**Table 2.** Apparent rate constants (k) for fucosylated vs. non-fucosylated glycans based on exponential saturation model fitting (n = 4; duplicate reactions × duplicate injections).

| Glycan Pair | Non-Fucosylated<br>$k$ ($\text{hr}^{-1}$) | Fucosylated<br>$k$ ($\text{hr}^{-1}$) | Fold Increase |
| --- | --- | --- | --- |
| G0 vs G0F | $2.39 \pm 0.05$ | $8.67 \pm 0.10$ | $\sim 3.6\times$ |
| G1' vs G1F' | $0.80 \pm 0.02$ | $3.28 \pm 0.03$ | $\sim 4.1\times$ |
| G1 vs G1F | $3.16 \pm 0.08$ | $3.69 \pm 0.09$ | $\sim 1.2\times$ |

**Table 2.** Apparent rate constants ( $k$ ) for fucosylated vs. non-fucosylated glycans based on exponential saturation model fitting ( $n = 4$ ; duplicate reactions × duplicate injections).
| Glycan Pair | Non-Fucosylated<br>$k$ (hr$^{-1}$) | Fucosylated<br>$k$ (hr$^{-1}$) | Fold Increase |
| --- | --- | --- | --- |
| G0 vs G0F | 2.39±0.05 | 8.67±0.10 | $\sim 3.6\times$ |
| G1' vs G1F' | 0.80±0.02 | 3.28±0.03 | $\sim 4.1\times$ |
| G1 vs G1F | 3.16±0.08 | 3.69±0.09 | $\sim 1.2\times$ |

The glycans G0 vs G0F and G1′ vs G1F′ showed a significant increase in their rate constants likely due to fucosylation of the glycan core. Although G1 vs G1F only showed a slight difference, with a 1.2-fold increase, the overall trend of fucosylated glycans is likely linked to a higher observed rate constant. Previous studies have shown that afucosylated therapeutic antibodies exhibit enhanced binding to their FcγRIIIa gamma receptor, which is expressed on immune cells, including natural killer (NK) cells. This stronger receptor binding enhances the ability to mediate antibody-dependent cellular toxicity (ADCC) compared to their fucosylated counterparts. However, fucosylated glycans are generally associated with reduced receptor binding due to steric hindrance between the Fcγ receptors and antibodies. This results in reduced ADCC, therefore decreasing therapeutic potency.^34^ The results of our study reveal a contrasting effect from a chemical-biology perspective, where the presence of core fucose is associated with increased procainamide labeling reaction efficiency.

One plausible explanation is that core or antennary fucosylation alters the conformation or flexibility of the glycan core, thereby increasing the fraction of the reducing end in an open chain (aldehyde) form accessible for reaction. Previous studies on fucose-containing oligosaccharides have shown that fucosylation can reduce conformational entropy (i.e. reduce flexibility) and favor a pre-organized or more rigid conformation, which lessens the entropic penalty for binding with lectins and possibly other reactive partners.^35^ Furthermore, the electronic nature of fucose (a 6-deoxy sugar) may slightly influence the electronic density at the reducing terminal N-acetylglucosamine by inductive effects, making the carbonyl carbon more electrophilic and reactive toward the amine moiety of procainamide. While direct studies of reductive amination kinetics in this context are scarce, analogous effects are seen in glycosyltransferase substrate specificity. Differential rates of transfer for fucosylated vs non-fucosylated acceptors are observed, suggesting that presence of a fucose moiety can promote or facilitate reactivity in neighbouring sugar residues.^36^ Microsolvation and steric effects are likely contributing as well where the fucosylation could alter local hydrophobicity or reduce hindrance around the reducing end, potentially improving access to labeling reagents. These structural micro-environments can accelerate the initial Schiff base formation, which is often the rate determining step in reductive amination. Such conformational and microenvironmental effects have also been implicated in lectin binding studies.^35^ The fundamental mechanism remains unclear. However, the factors involving improved reducing end accessibility, conformational pre-organization, and modest electronic effects may collectively contribute to the observed higher rate constants for fucosylated glycans during procainamide labeling. These findings prompt further research into how the fucose moiety can influence glycan labeling kinetics during reductive amination reactions.

### Determination of reaction time needed for efficient procainamide labeling of N-glycans

To quantitatively assess each glycan type labeling efficiency, the time required to reach 95% of the maximum fluorescence signal (A_max_) was calculated from the exponential saturation model, based on the relation 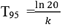 . As a model-based threshold to indicate the completeness of glycan labeling, the T_95_ benchmark was selected. The T□□ parameter is widely used in kinetic analysis for enzyme assays, reaction mechanisms, and labeling workflows. Therefore, it was chosen as the measure of analytical reliability to evaluate labeling efficiency.^37^

Figure 5. summarizes the T_95_ values for all nine glycans detected from TmAb, and the blue bar plot indicates that majority of the released glycans reached 95% of maximum labeling signal within 1 h. Glycans with rapid labeling reaction, such as G0F, Man5, and G0F-N, reached 95% of A_max_ in less than 0.6 h. In contrast, G1′ had the slowest kinetics, taking approximately 3.8 h to reach saturation, while G2F and G0 took 1.2 to 1.3 h to reach T□□. These results led to a proposal of three incubation time windows to balance labeling completeness versus analytical speed. For time-sensitive workflows, a 1 h incubation time is sufficient to achieve 95% or more labeling for most N-glycans. An optimal balance between speed and labeling requires an incubation time of 1.5 h. In applications requiring maximum labeling, to detect all minor glycans such as G1’, an incubation period of 2 to 2.5 h produces nearly complete labeling. These recommended incubation times offer researchers with flexible guidelines for various analytical goals, ranging from high-throughput screening to robust structural glycan profiling for PAT-based N-glycan analysis.

**Figure 5.**
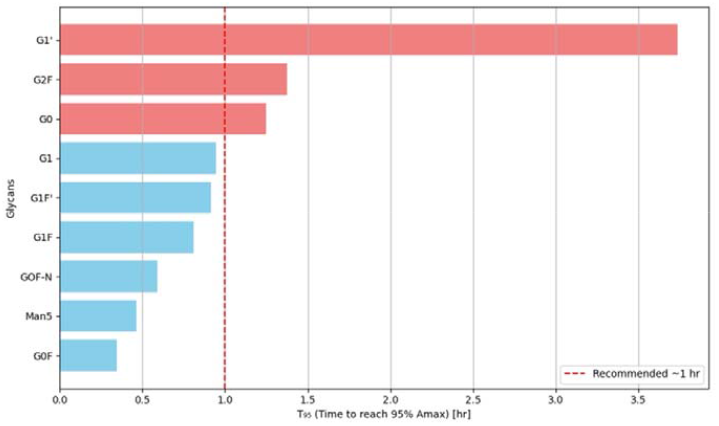
Comparison of T □ □ values, representing the estimated time required to achieve 95% labeling completion for each glycan, as determined from exponential saturation model fits.

## Conclusion

This study demonstrates that procainamide labeling of N-glycans by reductive amination can be significantly optimized for reaction time without compromising labeling efficiency. Most glycans reach near-complete labeling within one hour. The exponential saturation model fitted to the data generated in this study offers a straightforward and reliable approach to describing reductive amination kinetics, with rate parameters (such as A_max_ and k), that enable quantitative comparisons of labeling efficiencies across different N-glycan structures. Structural features of various N-glycans were seen to be impacting reductive amination rate and procainamide labeling efficiency. For example, fucosylated glycans were seen to label much faster than its afucosylated counterparts. Conversely, increased galactosylation likely causes steric hindrance to lower derivatization rates. These results enhance our understanding of N-glycan reactivity to reductive amination and support reducing procainamide incubation times to 1-1.5 h for higher-throughput applications relevant to real-time bioprocess monitoring. By defining kinetic thresholds, such as T_95_, this work also provides practical incubation time ranges suitable for various analytical needs, ranging from rapid high-throughput screening to detailed glycan composition profiling. Future research should systematically explore how branching patterns, terminal residues, and core modifications influence labeling efficiency to clarify some of the unresolved structure-kinetic relationships. Expanding this analysis by incorporating complementary structural methods, such as mass spectrometry, and more advanced kinetic models may provide deeper mechanistic insights. These efforts would enhance the predictive accuracy of kinetic models and facilitate the development of optimized labeling chemistries for advanced PAT-based glycomics for advanced continuous biomanufacturing of glycobiologics.

## Acknowledgements

This work was supported by awards #PC5.2-112 and #PC7.1-146 granted to Rutgers University under a Project Award Agreement from the National Institute for Innovation in Manufacturing Biopharmaceuticals (NIIMBL) and financial assistance award 70NANB21H086 from the US Department of Commerce, National Institute of Standards and Technology.

## Declaration of competing interest

The authors declare that they have no known competing financial interests or personal relationships that could have appeared to influence the work reported in this paper.

## Graphical Abstract

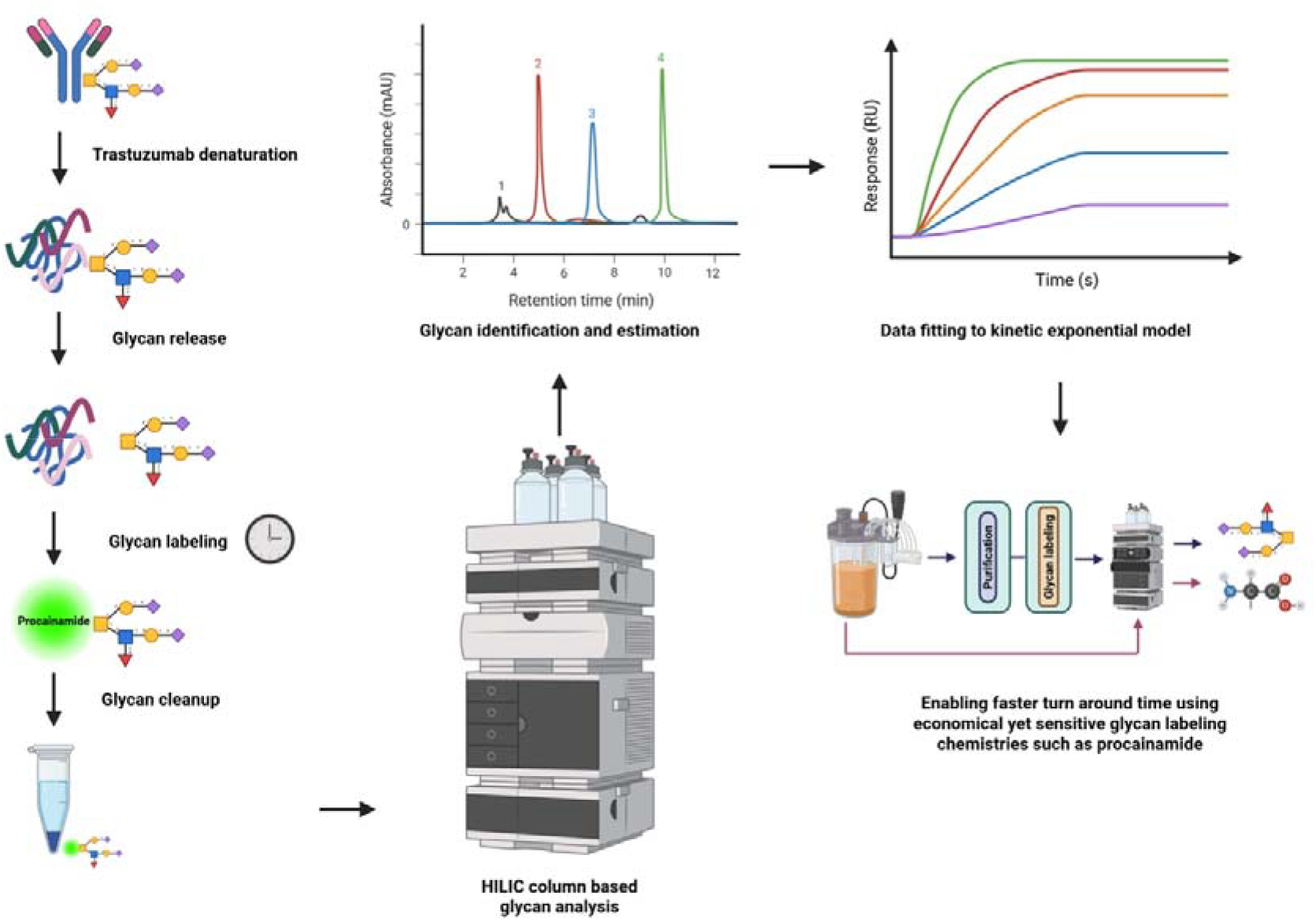

## Notes

### Competing Interest Statement

The authors have declared no competing interest.

