## Supplementary for "Reaction kinetics of procainamide dye derivatization of N-linked glycans to enable robust process analytical workflows for glycoprotein-based biologics manufacturing"

**Supplementary Table S1**. Glycan structures

| G0 | 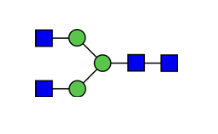 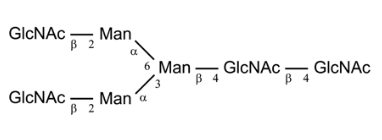 |
| --- | --- |
| G0F | 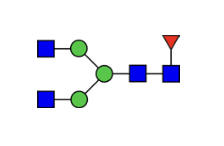 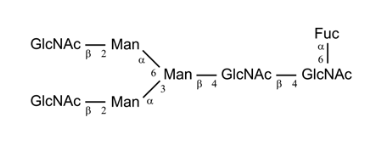 |
| G1 | 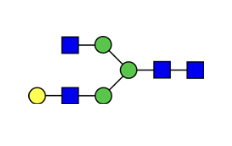 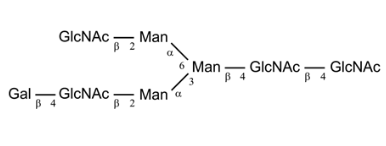 |
| G1’ | 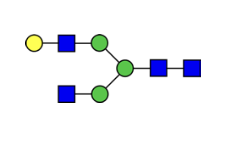 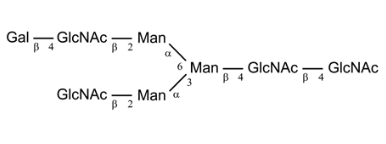 |
| G1F | 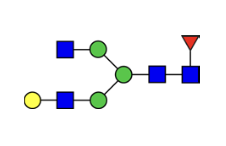 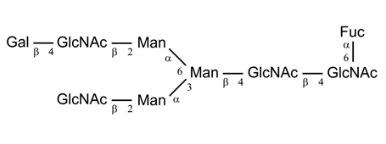 |
| G1F’ | 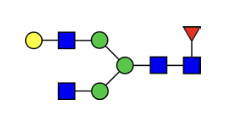 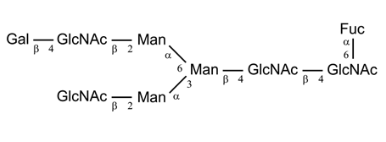 |
| G2F | 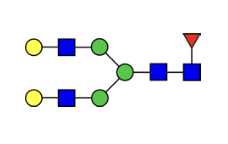 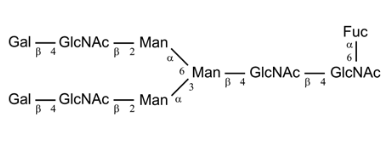 |
| G0F-N | 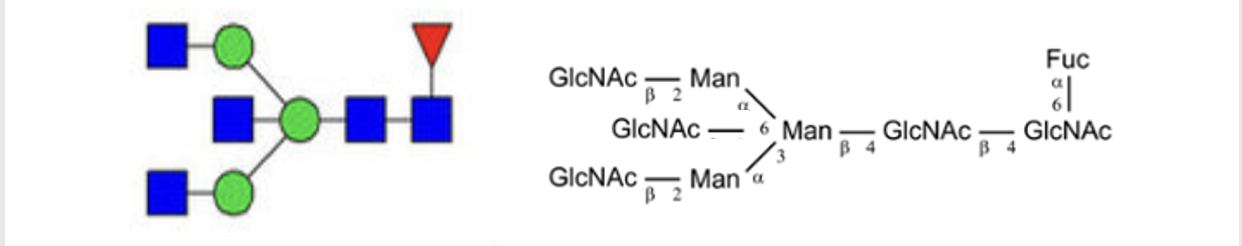 |

**Supplementary Figure S1**. Chromatogram of procainamide labeled glycans estimated for Trastuzumab biosimilar using HPLC.

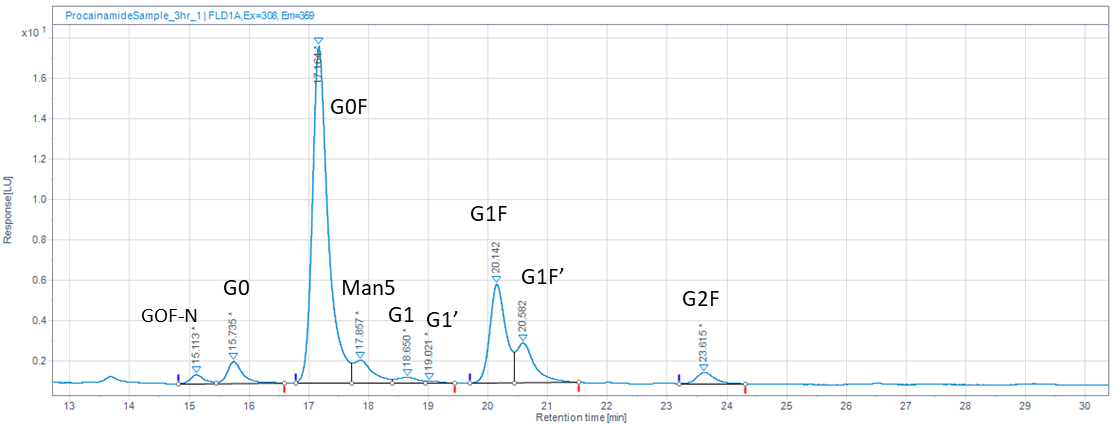

**Supplementary Table S2**. Estimation of area under the curve for each eluting procainamide labeled glycans of Trastuzumab biosimilar.

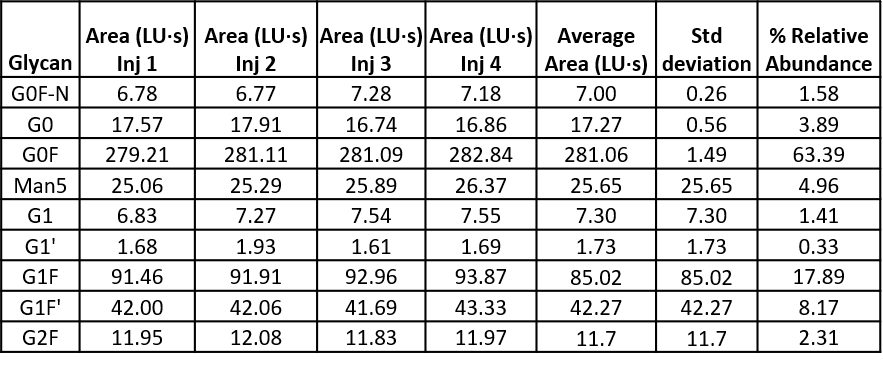

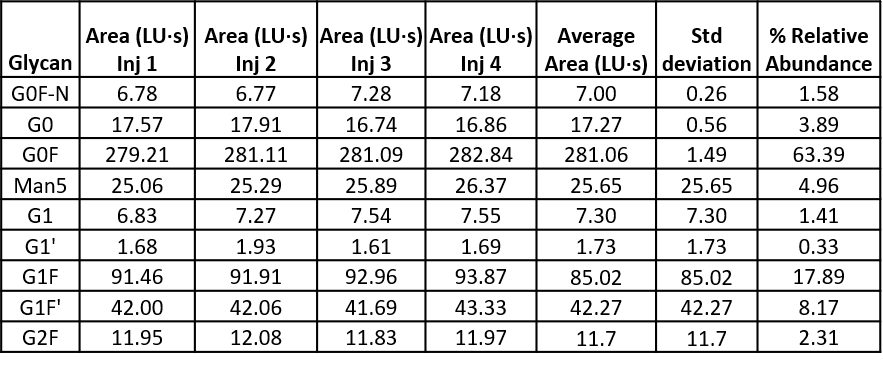
